# iSODA: A Comprehensive Tool for Integrative Omics Data Analysis in Single- and Multi-Omics Experiments

**DOI:** 10.1101/2024.08.02.605811

**Authors:** Damien Olivier-Jimenez, Rico J. E. Derks, Oscar Harari, Carlos Cruchaga, Muhammad Ali, Alessandro Ori, Domenico Di Fraia, Birol Cabukusta, Andy Henrie, Martin Giera, Yassene Mohammed

## Abstract

Omics technologies including genomics, proteomics, metabolomics, and lipidomics allow profound insights into health and disease. Thanks to plummeting costs of continuously evolving omics analytical platforms, research centers collect multi-omics data more routinely. They are, however, confronted with the lack of a versatile software solution to harmoniously analyze single-omics data and merge and interpret multi-omics data. We have developed iSODA, an interactive web-based application for the analysis of single-as well as multi-omics omics data. The software tool emphasizes intuitive, interactive visualizations designed for user-driven data exploration. Researchers can filter and normalize their datasets and access a variety of functions ranging from simple data visualization like volcano plots and PCA, to advanced functional analyses like enrichment analysis for proteomics and saturation analysis for lipidomics. For insights from integrated multi-omics, iSODA incorporates Multi-Omics Factor Analysis – MOFA, and Similarity Network Fusion – SNF. All results are presented in interactive plots with the possibility of downloading plots and associated data. The ability to adapt the imported data on-the-fly allows for tasks such as removal of outlier samples or failed features, various imputation strategies, or data normalization. The modular design allows for extensions with new analyses and plots. The software is accessible under http://isoda.online/.

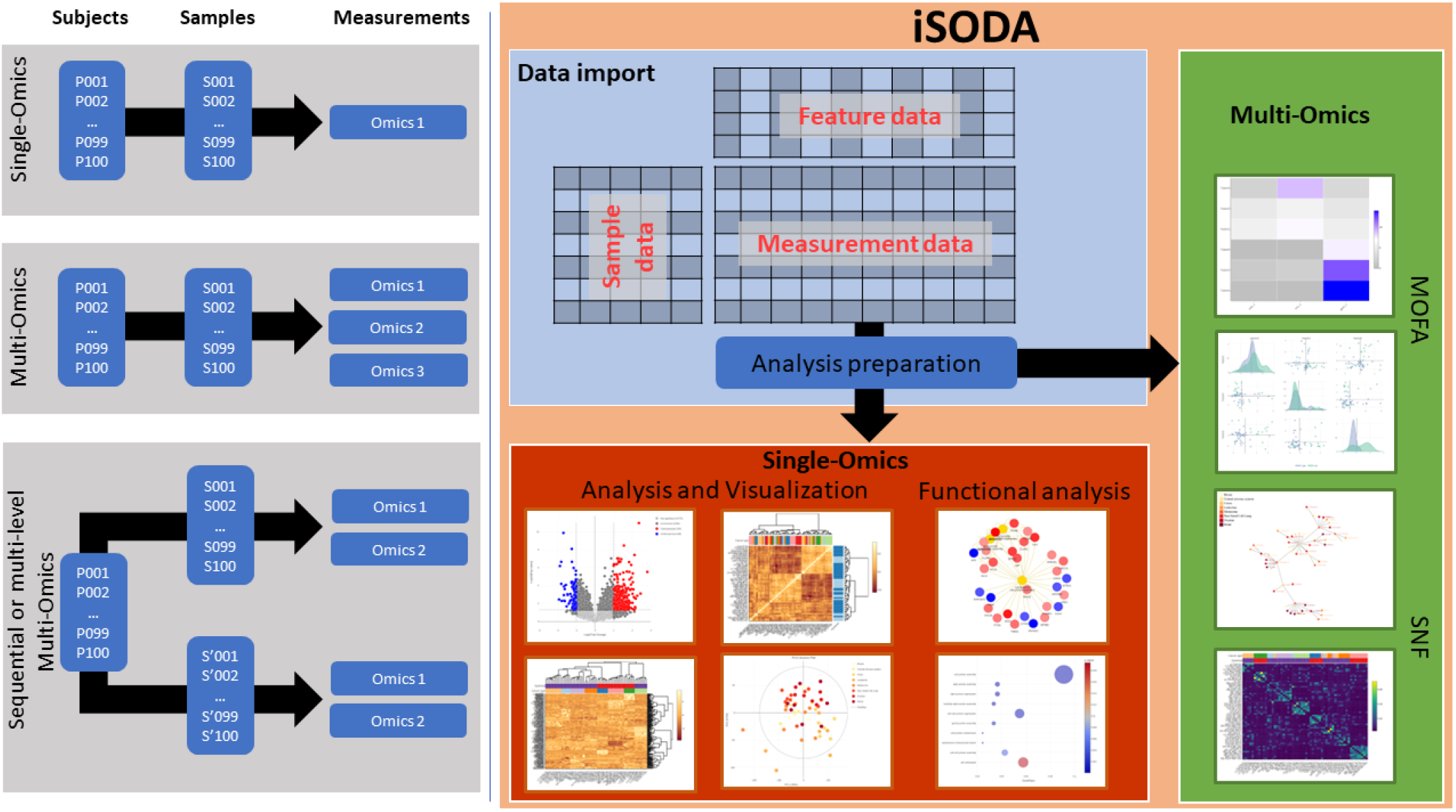

Graphical summary for iSODA showcasing some application examples, the data import, the single-omics and multi-omics modules.

## Introduction

The rapid advances in high-throughput omics resulted in an increase in data generation at lower costs, encompassing genomics, transcriptomics, proteomics, metabolomics, and beyond (1). These data help understanding biological processes, elucidating disease pathways, and advancing personalized medicine (2). Various tools and methods exist to navigate omics data — from statistical techniques pinpointing significant patterns to network-based approaches highlighting relationships within biological systems. Most tools, however, require either advanced knowledge of coding for data handling and/or dedicated to specific omics platform.

A few suites dedicated to analysis of (multi-)omics data have emerged over the past decade. Some focus on specific diseases or applications (3-8), while others adopt a general approach (9-16). Despite the long list of software tools (Table S1), a few notable implementations stand out for their interactive design and generality. MetaboAnalyst is a web-based application for metabolomics data analysis and interpretation (17). It currently allows streamlined analysis for both quantitative and untargeted metabolomics data. Perseus is specialized in interpreting protein quantification, interaction and post-translational modification data (18). It enables high-dimensional omics data analysis covering normalization, pattern recognition, time-series analysis, and classification. PaintOmics 4 maps acquired omics data on biological pathways (19) including KEGG (20) and Reactome (21). It supports pathways from KEGG, Reactome and MapMan. MergeOmics is pipeline for integrating datasets across omics (22). It leverages three functional analyses for disease association including Marker Set Enrichment Analysis (MSEA) to detect relevant processes, Meta-MSEA to examine the consistency between omics datasets, and Key Driver Analysis (KDA) to identify regulators of disease-associated pathways. OmicsAnalyst focuses on multi-omics analysis through three analysis tracks; correlation network analysis, cluster heatmap analysis, and dimension reduction analysis (10). Each of the tracks offers various algorithms to choose from. 3Omics generates inter-omics correlation networks with respect to experimental conditions for transcriptomics, proteomics and metabolomics (14). It includes coexpression analysis, phenotype analysis for transcriptomics and proteomics, KEGG pathway enrichment analysis for metabolomics and proteomics, and Gene Ontology enrichment analysis for proteomics and transcriptomics. In summary, great advances have been made towards convergent multi-omics data analysis. However, most tools take a deterministic approach to data analysis, making it a linear pipeline rather than exploration. The tools are either specialized for a specific omics, or dedicated to multi-omics, but not both. Static plotting seems also to be common in contrast to interactivity. All these aspects limit the users’ experience and ability to explore the data dynamically.

To that end, we have developed iSODA (integrated Simple Omics Data Analysis) as an interactive, expandable, and user-friendly software solution for single- and multi-omics data exploration. Our goal was to build a web-based platform that offers users a dynamic environment to process, study, and integrate their omics data; essentially to study multi-omics characterization experiments, on each omics layer individually, and side by side with multi-omics integration. Our goal was to transcend the linear approach to data analysis to enable a dynamic interactive exploration. We aimed to simplify the process for novice users without compromising the in-depth analysis for advanced users.

We demonstrate iSODA using two datasets. The first is a lipidomics library of 90 lipid transport protein knockouts studying the effect of these protein on the cell lipidome (23). In the second we use the multiomic characterization datasets of the NCI-60 cell lines (24), which were acquired in various laboratories. We discuss how to identify insights using the interactivity available in iSODA.

## Materials and Methods

### Software implementation

iSODA was developed in R – 4.4.0, and Shiny framework (25) for the graphical interface. We relied on shiny extensions including bs4Dash (26) and ShinyWidgets (27) for additional UI elements, shinymanager (28) for user authentication, shinyjs (29) and shinybrowser (30) for aspect ratio adjustment. The backend architecture is object-oriented implementing three R6 (31) classes; 1) the omics class for single omics experiments with methods for data import, processing, visualization, and result storage; 2) the MOFA class for Multi Omics Factor Analysis (32); and 3) the SNF class for Similarity Network Fusion integration (33). The MOFA and SNF classes contain the methods for importing data from the single omics classes, processing them accordingly, and visualizing the results. Omics data are imported into iSODA as tables in comma-separated (csv), semicolon-separated (csv2), tab-separated (tsv) or excel (xlsx) formats. We used plotly (34) for interactive plots, and visNework (35) for visualizing networks. Finally, the heatmaply package was used to visualize heatmaps (36). For all plots, a various color palettes are available from RColorBrewer, viridisLite and grDevices packages.

The various analyses rely on the stats core package including: Distance calculations – Euclidean, maximum, Manhattan, Canberra, binary, and Minkowski. Hierarchical clustering methods – ward.D, ward.D2, single, complete, average, McQuitty, median, and centroid. Pearson’s and Spearman’s correlation coefficients for correlation, Student’s t-test and Mann–Whitney–Wilcoxon for statistical tests, and p-value adjustment methods include Bonferroni (“bonferroni”), Holm (“holm”), Hochberg (“hochberg”), Hommel (“hommel”), Benjamini & Hochberg (“BH”), and Benjamini & Yekutieli correction (“BY”). Principal Component Analysis – PCA using pcaMethods package (37). Advanced regression analysis and feature selection was achieved using Lasso and Elastic-Net Regularized Generalized Linear Models using the glmnet package (38). For functional analyses we used ClusterProfiler and Enrichplot packages for enrichment and over-representation analyses (39-41). Gene and protein annotations are provided using the Org.Hs.eg.db package (42), and users can also upload their own annotation for any omics for functional analyses.

### LTP KO lipidomics library

The Lipid Transfer Protein Knockout (LTP KO) library was produced by our group (23) to investigate the disturbances caused by the loss of lipid transfer proteins (LTPs). Briefly, the MelJuSo cell line (human melanoma) was used and 90 LTP gene knockouts experiments were conducted using CRISPR/Cas9 technology (Table S2). These 90 genes are from 11 families determined by the LTP domains: OSBP (n=30), START (n=34), PITP (n=16), GLTP (n=11), CRAL-TRIO (n=71), SMP (n=15), VPS13 (n=11), NPC1 NTD (n=6), ML (n=9), SCP2 (n=12) and ASTER/VAST (n=8). Additionally, six non-targeting controls were included. These samples were analyzed for their lipidomics content in the original study. An extensive description of the samples and measurement is available in the original work (23).

### NCI-60 cancer cell lines multi-omics dataset

The 60 Human Tumor Cell Lines used by National Cancer Institute for over 20 years as a screen to identify and characterize novel compounds for growth inhibition or killing of tumor cell lines (43). The screen utilizes 60 different human tumor cell lines, representing leukemia (LE, n=6), melanoma (ME, n=10), and cancers of breast (BR, n=5), central nervous system/brain (CNS, n=6), colon (CO, n=7), non-small cell lung (LC, n=9), ovarian (OV, n=7), prostate (PR, n=2) and renal (RE, n=8). Table S3 lists the individual cell lines. CellMiner is an online database hosting the multi-omics characterization data on the NCI-60 cell lines (24,44). We used three characterization datasets representing genomics (Illumina 450K methylation, gene average), transcriptomics (5 Platform Gene Transcript, averaged intensities) and proteomics (SWATH mass spectrometry, protein). A complete description of the sample measurement and data acquisition are provided in the original publication (24,44).

### Data preparation

All datasets went through a reformatting step to ensure compatibility with iSODA simple format. Two essential tables were generated for all datasets, i.e. data table and sample annotation table. In addition, when available, a feature annotation table was generated and used. iSODA requires the first two tables for analysis and visualization, while the feature annotation is required to map feature metadata onto the visualizations and for the functional analyses, i.e. enrichment and over representation analysis. In case of gene associated measurement (genomics, transcriptomics, proteomics), an internal annotation based on Gene Ontology will also be available.

## Results

We give next an overview of iSODA and we follow with a discussion of two use-cases.

### iSODA – integrated Simple Omics Data Analysis

Each omics layer has a dedicated interface with data upload, visualization, and functional analysis tabs. The multi-omics modules allow selecting all or some of the data in the single-omics experiments for integration (Figure S1). We describe here the functionalities briefly, and include a comprehensive description in the supplementary materials.

### Single-Omics data analysis

Users upload their data via three subtabs for: *Sample annotations, Measurement data*, and *Feature annotations*. ***Data pre-analysis*** includes filtering, normalization (scaling, total normalization, standardization), imputation (minimum, median, mean, maximum), and batch effect correction using ComBat (45). In ***Interactive visualization*** up to four plots can be displayed simultaneously, and all can be interacted with by zooming and hovering. Various display and analysis parameters can be adjusted for each plot from the parameter sidebars. ***Dendrogram*** *p*rovides a rapid assessment of the similarity between samples and sample groups via hierarchical clustering. ***Volcano Plot*** visualizes features that differentiate between two groups using adjustable statistical tests and fold change thresholds. Additional information regarding the features can be mapped onto this plot, for instance, Gene Ontology terms (genes/proteins) or chemical classes (lipidomics). The ***heatmap*** can be used to cluster samples and features allowing for the identification of sample groups and features groups. Outlier samples or failed measurements can be spotted here. Uploaded sample annotations can also be mapped, and using LASSO and Elastic-Net Regularized Generalized Linear Models allow feature selection for segregating sample groups. This provides a clear view of how sample groups correlate with specific features. ***Principal Component Analysis*** (PCA) visualizes samples and features in a reduced dimensional space maximizing variance highlighting unsupervised groupings. Plots for explained variance, scores, and loadings are available. Annotations enhance interpretation and Lasso helps identify key features driving trends. ***Sample Correlation Heatmap*** provides a visual representation of the correlations between samples, allowing the user to spot grouping and batch effects. ***Feature Correlation Heatmap*** provides similar information but applied to the features. All plots can be downloaded in several formats (png, svg, jpg, webp).

iSODA allows the implementation of omics specific analyses. ***Class Distribution*** is a lipidomics specific visualization that provides a summary of the mean lipid class concentrations for all sample groups. ***Class Comparison*** is a stratified version of the class distribution, allowing a better assessment of the minute group concentration variations. ***Double Bonds Plot*** complements the volcano plot by focusing on the differences in double bounds in lipid classes between two groups.

***Functional analysis*** implements Enrichment Analysis (EA) and Over-Representation Analysis (ORA). EA is based on modified K-S statistics as implemented originally by Geneset Enrichment Analysis (GSEA) (46). Over-representation Analysis (ORA) uses Fisher exact test to identify if a feature annotation is over-represented in a list of pre-selected features of interest compared to random chance (47). The user can upload their own feature annotations and perform EA and ORA. Gene Ontology terms are made readily available for corresponding omics, i.e. genomics, transcriptomics and proteomics. ***Dot Plot and Bar plot*** display the significant feature sets resulting from the analysis according to the test’s p-values. ***Ridge Plot*** provides density curves of features (genes, proteins, etc.) associated with each enriched term/function. ***CNET Plot*** is an interactive network representation of measured features and associated enriched terms. The interactive network enables a physics mode in which nodes (enriched terms) are pulled together or repelled based on their association with measured features (genes, proteins, etc.). This would cluster terms together in a data-driven way based on features and annotations. ***eMap Plot*** is an enrichment map plot similar to CNET plot, but specifically designed to visualize larger numbers of feature sets.

### Multi-omics integration

The integration provides a holistic view allowing to pinpoint patterns that are not readily observable in each single-omics alone. In the current implementation, iSODA offers two multi-omics integration strategies, i.e. Multi-Omics Factor Analysis (MOFA) and Similarity Network Fusion (SNF).

***MOFA*** is an unsupervised integration designed to reduce the complexity of large-scale omics datasets into a manageable number of latent factors (32). These factors capture the underlying sources of variation across the omics datasets, providing insights into driving biological processes. In a way, MOFA’s factors are similar to PCA components in one omics layer. ***Explained Variance Plot*** provides an overview of the contribution of each omics dataset to each factor. It allows identifying factors dominated by a specific omics as well as those that are shared across multiple omics. ***Factor Plot*** summarizes the sample factor weights and assesses their potential to explain group differences. ***Combined Factors Plot*** is a scatter plot of sample factor weights from two factors allowing to explain dependency/independence between factors. ***Feature Weights Plot*** shows the contributions of individual measured features to a selected factor within an omics dataset. ***Feature Top Weights Plot*** focuses specifically on the highest-contributing features in each factor and displays these as lollipop plot. ***MOFA Heatmap*** shows the factors’ top contributing features versus the samples enabling a visual inspection of how certain sample groups can be clustered by relevant features in different omics. ***Scatter Plot*** allows examination of how the measured top-weight features correlate with sample weights.

***SNF*** constructs individual networks for each dataset, each representing the similarity between samples. These networks are then fused into a single network through an iterative process capturing both the shared and unique characteristics of each dataset (33). ***Similarity Heatmap*** illustrates the impact of clustering on omics datasets separately. The x- and y-axes display samples organized by cluster groups, while the cell values indicate similarity (SNF affinity) levels. Cluster assignments can be used to evaluate alignment with sample grouping. ***Fusion Heatmap*** organizes samples according to their cluster based on the integrated network. ***Similarity Network*** visualizes the affinity matrix for single-omics datasets as an interactive network, and ***Similarity Fusion Network*** represents the fused multi-omics sample clustering as an interactive network. By hovering over an edge between two sample nodes in the network, the user can display information on the evidence from the data.

### Lipidomics of lipid transfer protein knock-outs

Lipid transfer proteins (LTP) are classified into families based on protein domains suggesting similarities in function (48,49). There are currently 348 proteins annotated as lipid transport proteins (50). To systematically study and highlight the function of various LTPs, we characterized 90 intracellular LTP KO based on MelJuSo cell line using targeted quantitative lipidomics with internal standards (supplementary data Table S2, Figure S2) (23). Using iSODA we imputed missing values with median and removed lipid species below two times the average blank signals; 752 lipid species out of the measured 1109 remained and were used for further analysis.

First, we attempted to identify any association between the LTP families and observed lipidomics phenotype. The protein family is based on functional domain similarity, and suggests that similar functions would be reflected in respective lipid cargo giving rise to similar patterns in lipidomics, underlining associations between LTP families and their role in lipid metabolism. Our data revealed no such associations (Figure 1A). Considering that a protein domain might interact/affect specific lipid species, therefore the effect might only show on a subset of measured lipids, we performed a multinomial LASSO regression to extract any discriminatory lipid species between families. This also led to no observable discrimination (Figure 1A). It can hence be argued that protein domains and their similarities are not necessarily proxies for the actual function. Examination on a gene-by-gene basis seemed essential to identify the effects of a specific knock-out on the lipidome (Table S4). We highlight here a few examples.

**Figure 1.**
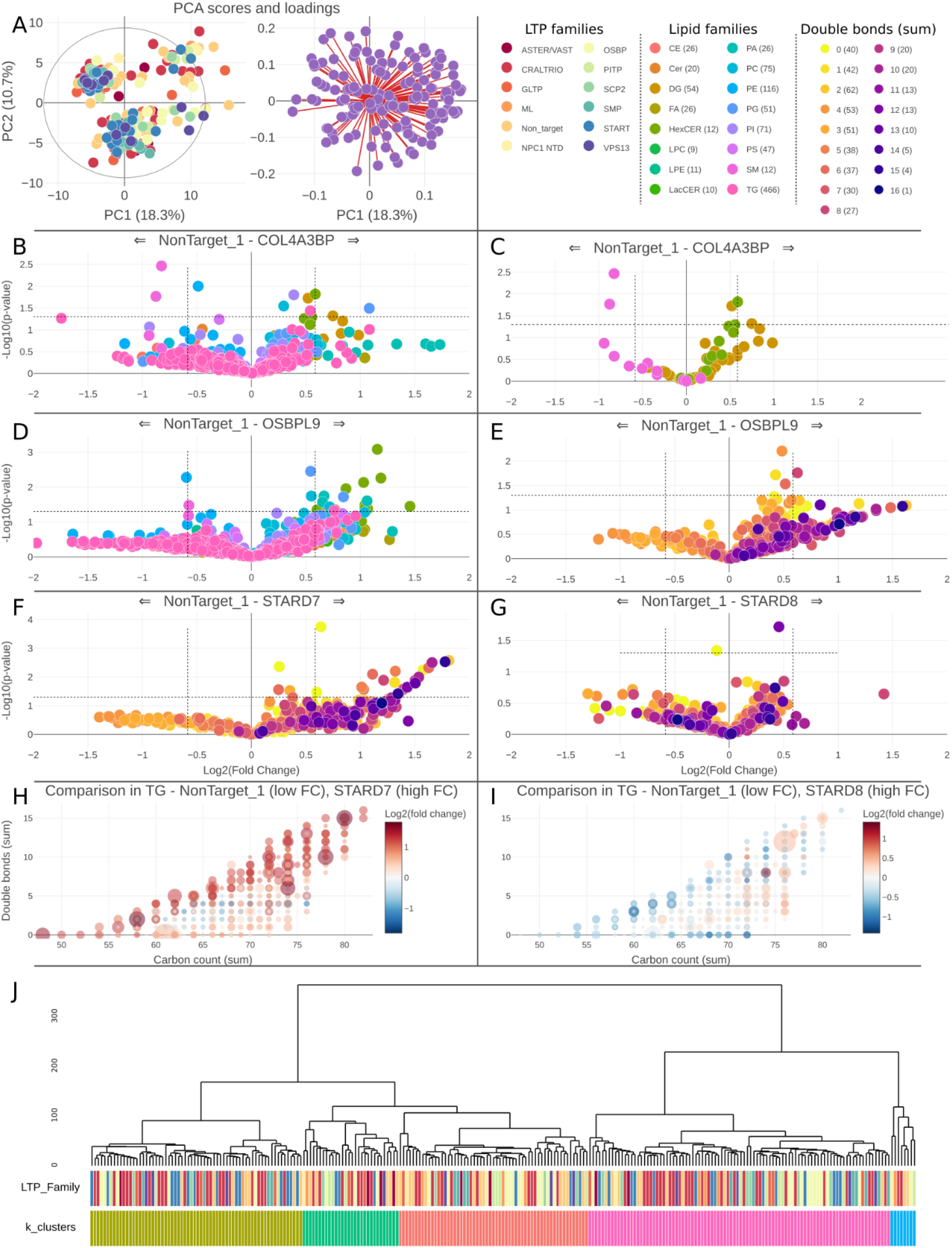
Lipidomics analysis of the LTP KO dataset. (A) PCA with scores and loadings of all KO colored according to family. (B) Volcano plot comparing COL4A3BP KO to non-target displaying lipid classes, and in (C) selected lipid classes that show trends are displayed. (D) Comparing OSBPL9 to non-target showing no visible trend for any lipid class, and (E) with only TGs colored by the total double bond count showing a clear trend. In (F) and (G) as well as (H) and (I) TGs in StarD7 and StarD8 were compared to non-target showing different trends regarding the double bond in the volcano plots (F, G), and in the detailed double bond plots (H, I). (J) A dendrogram of all LTP KO library using all lipid species measured to determine the distances showing no obvious alignment with the LTP family. Samples could however be clustered according to 5 main groups. Total normalized tables were used in all cases, lasso was applied to PCA based on the LTP family, alpha=0.8.

COL4A3BP, also known as Ceramide Transfer Protein (CERT), is a gene that encodes an LTP primarily transferring ceramides between the endoplasmic reticulum and Golgi apparatus. Ceramides are key components of cell membranes and play a crucial role in cellular signaling and apoptosis. Dysregulation of ceramide metabolism has been linked to various cancers. Moreover, given the importance of sphingolipids in the nervous system, dysfunction of COL4A3BP in known to be implicated in neurological diseases (51). When comparing the COL4A3BP KO to control (Figure 1B) a few significant lipid species can be detected. Mapping lipid class on the volcano plot revealed that DGs and LacCer species were present at higher concentrations while lipids from the SM class were at lower concentrations (Figure 1C). Although not at a significant level with FC of 1.5 (corresponding to logFC cutoff 0.585) a trend can be seen given all measured LacCer and SM species.

Next, we investigated Oxysterol Binding Protein-Like 9, OSBPL9, which is part of the oxysterol-binding protein (OSBP) family. It contains a pleckstrin homology domain, which facilitates binding to phosphoinositides, and a conserved oxysterol-binding domain that binds sterols and other lipids, and therefore is believed to binds and transports predominantly sterols and phosphoinositides within cells. OSBPL9 plays a crucial role in maintaining cellular lipid homeostasis and is involved in various cell signaling pathways regulating growth, survival, and metabolism. Disruptions in OSBPL9 may contribute to neurological disorders given how critical lipid homeostasis for neural functions is. Several lipid classes show overproduction in OSBPL9 knockout compared to control, notably Hexosylceramides (HexCer) with multiple significant species (Figure 1D). Moreover, when focusing specifically on Triglycerides (TG) and investigating the double bonds, additional patterns emerge with count of double bonds higher than 5 being gradually overexpressed in the knockouts (Figure 1E).

StarD7 and StarD8 belong to the StAR-related lipid transfer (START) domain family, known for their role in lipid binding and transfer. StarD7 is specifically involved in the intracellular transport of phospholipids and particularly the transfer of phosphatidylcholine to mitochondria, which is crucial for maintaining mitochondrial membrane integrity and function. StarD8, also known as DLC-3 (Deleted in Liver Cancer-3), is involved in lipid binding and transport, and plays a role in cellular signaling pathways by interacting with phosphoinositides. StarD8 is also known to regulate the activity of Rho family GTPases, which are critical for cytoskeletal dynamics, cell movement, and growth. Comparing the effects of the StarD7 and StarD8 knockouts on the cell lipidome shows that they have opposite effects within a single lipid class. Absence of StarD7 affects TG saturation, while StarD8 absence does not (Figure 1F, G). This effect can also be seen in more detail when using the double bond plots, specifically showing opposite effects for many of the lipid species examined (Figure 1H, I).

### Multi-omics characterization and integration of the NCI-60 cancer cell lines

The three omics datasets covered epigenomics, transcriptomics and proteomics of the NCI-60 cell lines. For epigenomics DNA-methylation levels were measured, for transcriptomics gene transcripts averages between 5 platform were used, and for proteomics SWATH (data independent acquisition) was performed on NCI-60 cell lines. After inspection in iSODA, prostate cancer with only two cell lines was underrepresented and therefore removed. Two other samples were excluded, i.e. ME:MDA-N for having no data in proteomics and CNS:SF-539 for having too many missing values in transcriptomics. The remaining cell lines covered 8 cancers; leukemia, melanoma, breast, central nervous system/brain, colon, non-small cell lung, ovarian, and renal.

### Single-omics analyses

The sample correlation heatmap (Figure 2A-C) shows that the cell lines clusters follow the cancer type in transcriptomics (Figure 2B), but not on proteomics and DNA-methylation level (Figure 2A, 2C). A closer look shows the epithelial nature of the cells, i.e. carcinoma or not, is a main driver of the grouping. This is most apparent in transcriptomics, which seems to carry most embedded patterns regarding cancer and epithelial nature. PCA score plots of both cancer and epithelial nature show similar trends (Figure 2D-F). Here, the non-epithelial cell lines form two groups with leukemia on top-right, and melanoma together with CNS at bottom-left (Figure 2E). Epithelial cell lines show distinction between colon on one side, and lung, ovarian, and renal cancer on the other side. A regression analysis to select discriminating features can be applied with provided sample annotations. When using cancer types as main grouping endpoint, various features in each of the three omics were identified (Figure 2J-L). Interestingly, the epithelial nature of cell lines still drove the clustering indicating an underlying strong similarity between cell lines based on origin. A similar analysis according to the epithelial nature of the cell lines was also performed (Figure S3).

**Figure 2.**
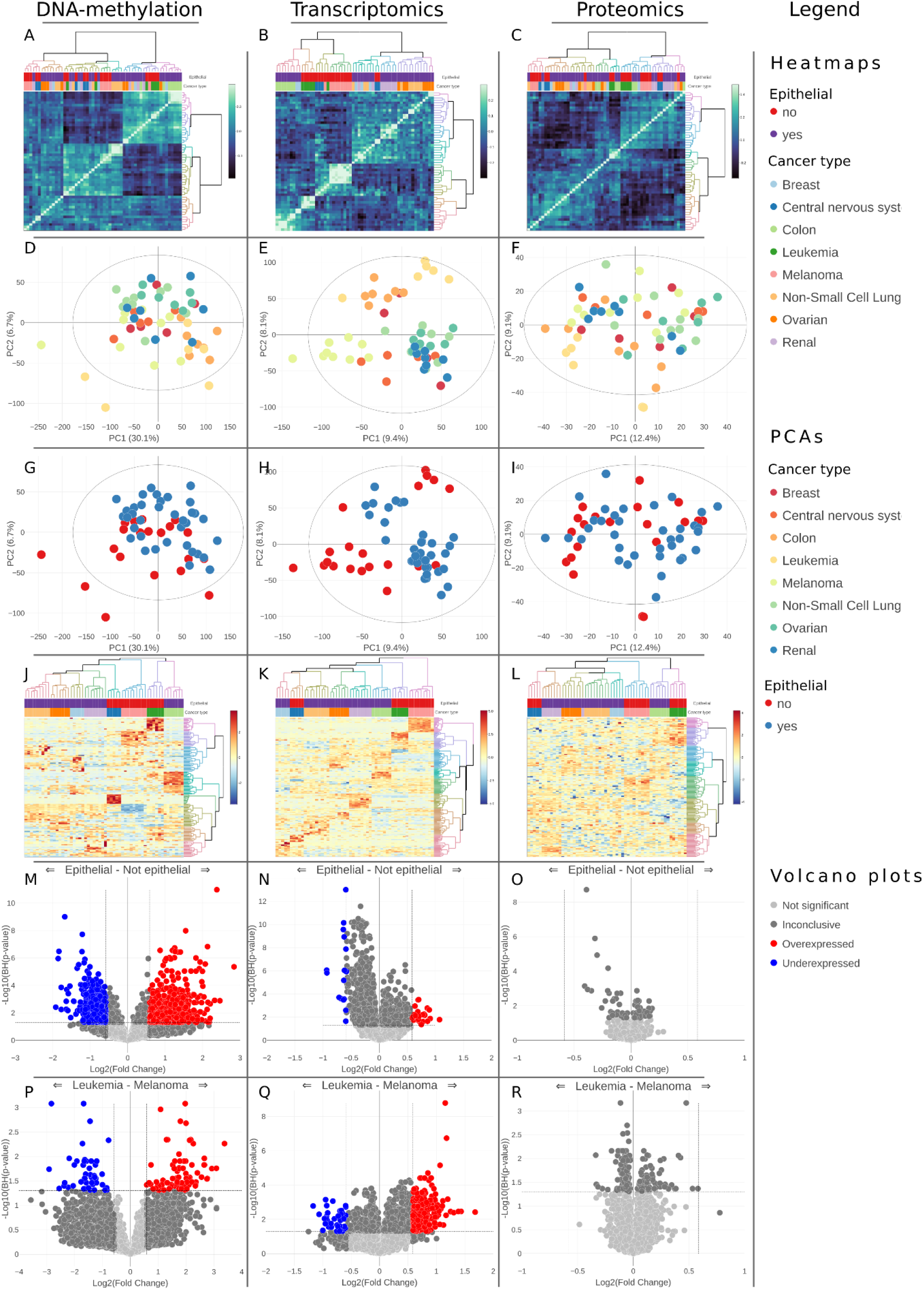
Single-omics data analysis of the NCI-60 DNA-methylation, transcriptomics, and proteomics characterization data. A-C correlation heatmaps showing similar clusters based on Pearson’s correlation. D-I represent PCA score plots (z-scored total normalized data, the nipals PCA method) with D-F are colored by cancer type and G-I by epithelial nature. J-L heatmaps with feature selection to discriminate cancer type for the three omics (z-scored total normalized data, LASSO alpha set to 0.8, Euclidean distance, ward.D2 clustering). Annotations mapped on top included cancer type and epithelial nature of the tissue. k=8 clusters was set and is rendered in the colored dendrogram. M-O volcano plots comparing cell lines according to their epithelial nature, and P-R comparing leukemia to melanoma samples (total normalized data, t-test, FC using mean, B-H p-value adjustment, p-value threshold was set to 0.05, FC threshold was set to 1.5).

To identify statistically significant features, we used volcano plots, and epithelial and non-epithelial cell lines were compared in all three omics. Many discriminating genes were identified in DNA-methylation data, with less in transcriptomics, and no significant proteins were found (Figure 2M-O). While the multivariate analysis using regression was able to identify features for such discrimination, the univariate analysis using statistical testing and fold change was less effective, save for DNA-methylation. These results appear to be intertwining; the epithelial nature of cells seems to be regulated upstream, hence DNA-methylation data shows discriminating features in volcano plots; however, the effect on the phenotype is more subtle appearing only in the multivariate analysis.

We also compared various cell lines in pairs, Figure 2P-R shows leukemia and melanoma, both non-epithelial. Although no proteins were significantly over/underrepresented, various transcripts and methylated genes were. The possibility to perform these analyses and interact with the results on the screen in a few clicks enables streamlined analysis and quick contrasting towards trends identification.

### Functional analysis

In addition to accepting user’s custom feature annotations for functional analyses, iSODA annotates genomics, transcriptomics and proteomics with Gene Ontology. From the various possible comparisons, we highlight the results of the enrichment analysis comparing melanoma to leukemia (Figure 3A-F). Interestingly, functions related to immune response were upregulated in the DNA-methylation, and downregulated in the transcriptomics data. This aligns with the general understanding that DNA methylation regulates gene transcription through repression. Here, we observe this repression on the functional level rather than on a specific gene. Various functions that exhibit opposite regulation in the DNA methylation versus transcriptomics were associated with different immune system pathways. Further, methylation of genes associated with synapses and morphogenesis were reduced in melanoma, while genes associated with pigmentation and cell junction were upregulated in melanoma. Over representation analysis of the differentiated gene methylation between epithelial and non-epithelial cell lines resulted in functions grouped into two clusters; associated with epidermis development and cell junction assembly (Figure 3G-H). The functional analyses demonstrate how different omics complement each other in describing cell biology. To achieve a more holistic view, we next describe omics integration.

**Figure 3.**
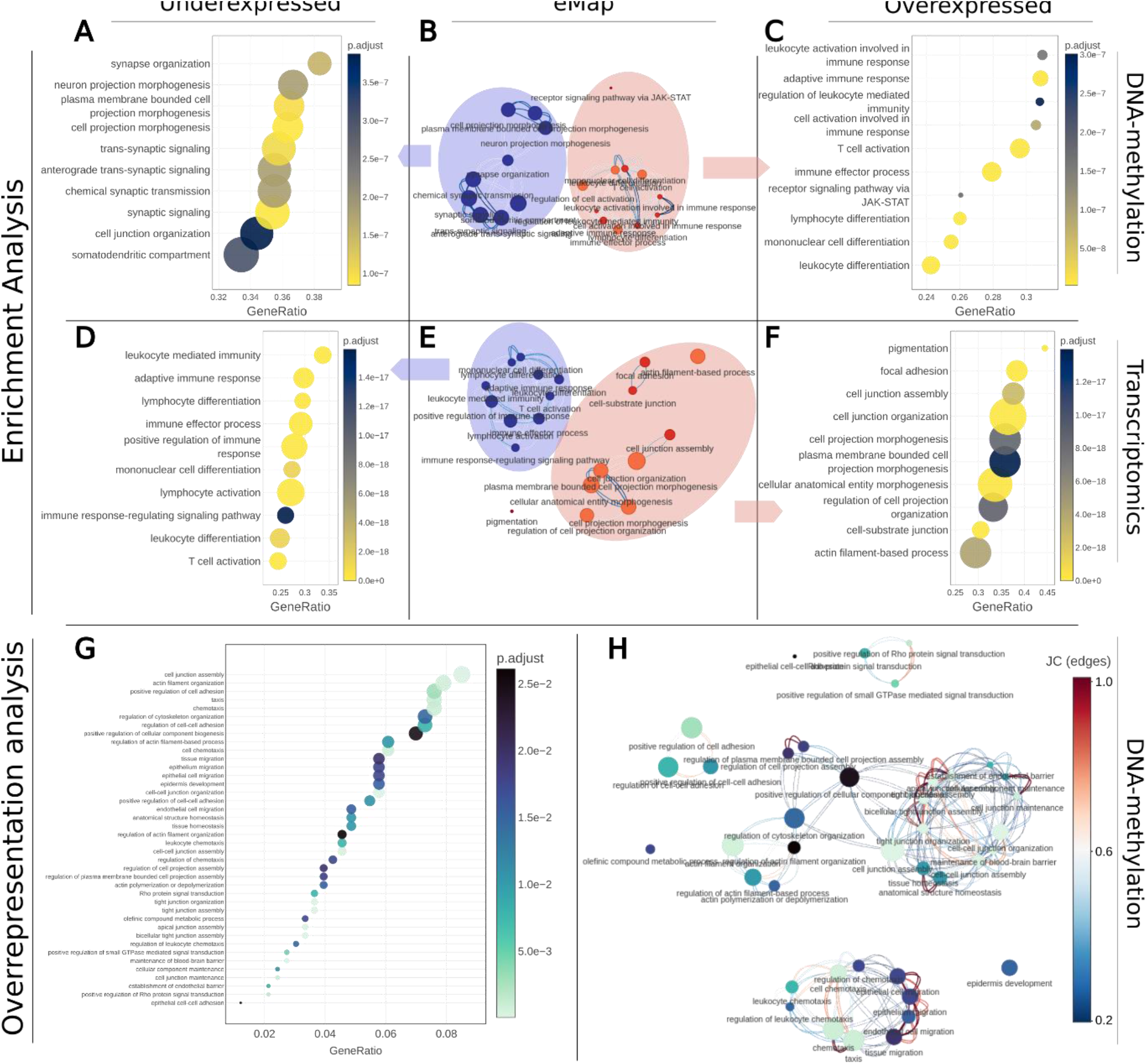
Functional analyses in NCI-60 omics characterization data. A-F show functional analyses with results for DNA-methylation and transcriptomics data comparing leukemia and melanoma cell lines. Feature set adjustment was set to BH, p-value=0.05, set minimum and maximum size were 3 and 800 respectively. A and C are dot plots and B is eMap from the DNA-methylation data, D and F along with E are for transcriptomics. In dot plots top 10 enriched terms were selected, in eMaps top 20, colored by NES values with blue for negative and red for positive, node size scaled to gene count, similarity score was JC, score threshold was 0.2. For each omics the suppressed (A, D) and activated (C, F) biological processes and molecular functions are shown. G and H show over representation analysis comparing epithelial and non-epithelial cell lines with the DNA-methylation data, using the under- and overexpressed features. Over- and underexpressed DNA-methylated genes were determined by the volcano plot analysis and imported into functional analysis using the built-in save tool feature. H shows top 20 annotations. G displaying top 20, node coloring reflects adjusted p-value, their size is scaled to gene counts, JC similarity score used with threshold set to 0.2, edges colored according to JC score.

### Similarity Network Fusion of the NCI-60 cell lines

SNF finds similarities between samples using spectral clustering (33). For comparison we show single-omics and multi-omics clustering (Figure 4). Before fusion, leukemia cell lines are barely clustered together, the renal cell lines produce compact clusters on DNA-methylation and transcriptomics, and the melanoma cell lines cluster generally consistently in all three datasets. When applying SNF, the integrated view better captures the nature of cell lines by clustering these – to a good degree – into their original cancer (Figure 4G, H). Interestingly, a data-driven k=8 clustering shows a better agreement with the original cancer type after SNF compared to each single omics alone. While SNF indicates integration enhances clustering, we turn to MOFA to study features driving the similarities.

**Figure 4.**
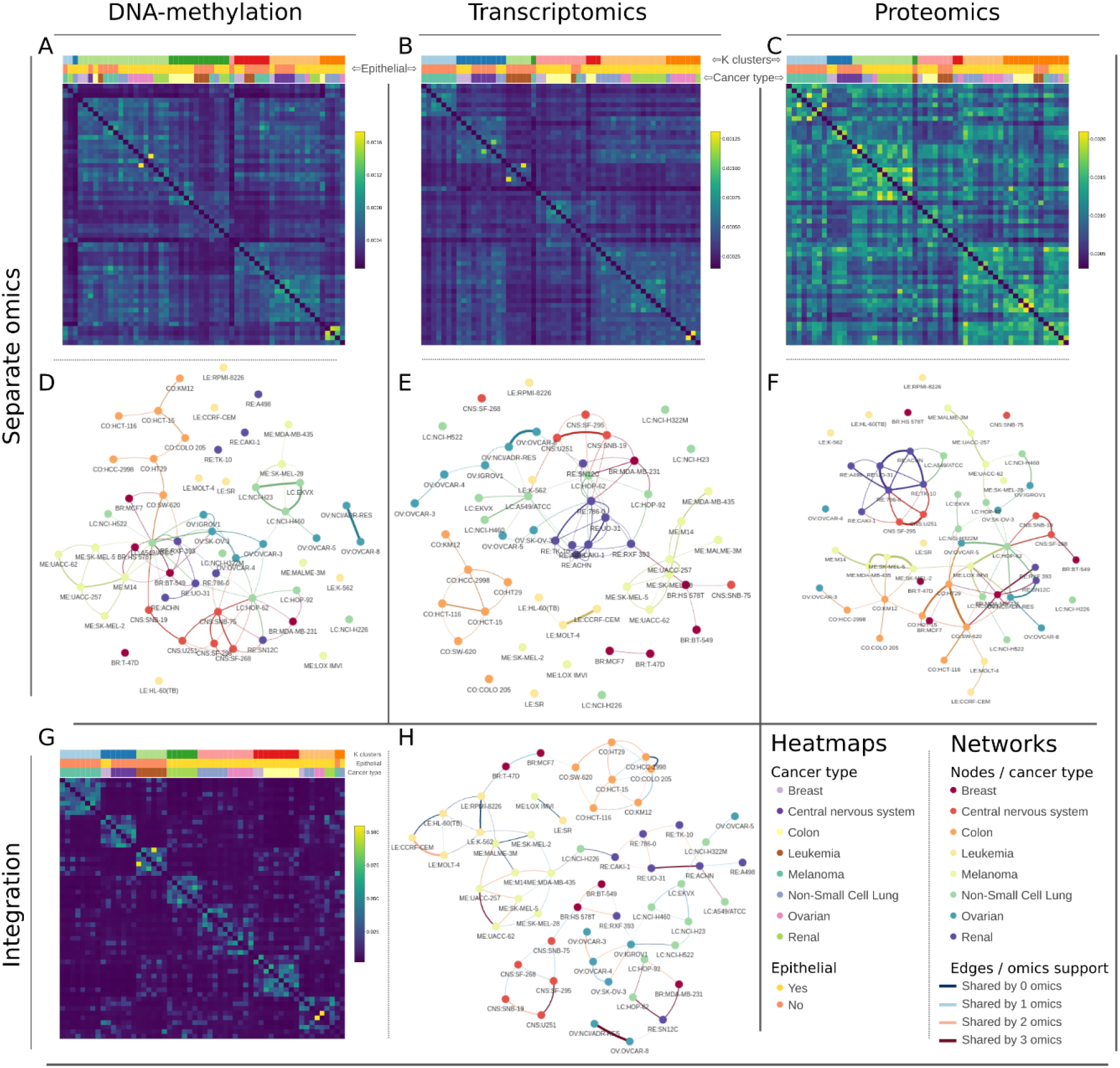
Similarity Network Fusion on NCI-60 cell lines. A-C are similarity heatmaps and D-F are the network of the three omics individually. G shows the fusion heatmap and network results. k-nearest neighbors method was set to 5, sigma to 0.5, using an Euclidean distance, and setting k clusters to 8. The heatmaps were mapped with k clusters, cancer type and epithelial status, while the networks were colored by cancer type and 5% of the top scoring edges were kept.

### Multi-Omics Factor Analysis of the NCI-60 cell lines

We set MOFA to generate 10 factors; however, the exact number of factors can vary according to the integration goal. For example, up to 10 factors are recommended to find measured features that drive the variance in the different omics layers, however, finding outlier samples requires more factors. Additional details can be found in the original MOFA documentations (32). The variance plot (Figure 5A) shows the explained variance associated with each factor and omics. The factor plot shows the samples’ factor weights and allows coloring according to the different sample annotations. When projecting the epithelial nature of the cell lines, there was no clear separation across all 10 factors. When projecting cancer type a few factors stood out, especially factors 1, 3, and 6 separating colon, leukemia and melanoma from the rest of the cell lines (Figure 5B-E). For an in-depth view of the integration results and how they relate to the data measured, the top contributing features for a factor can be displayed (Figure 5F-H) along with the measured values and associated sample weight (Figure 6I-K). Heatmaps of these top features and how they contribute to discrimination in each omics dataset separately can also be viewed and studied (Figure S4).

**Figure 5.**
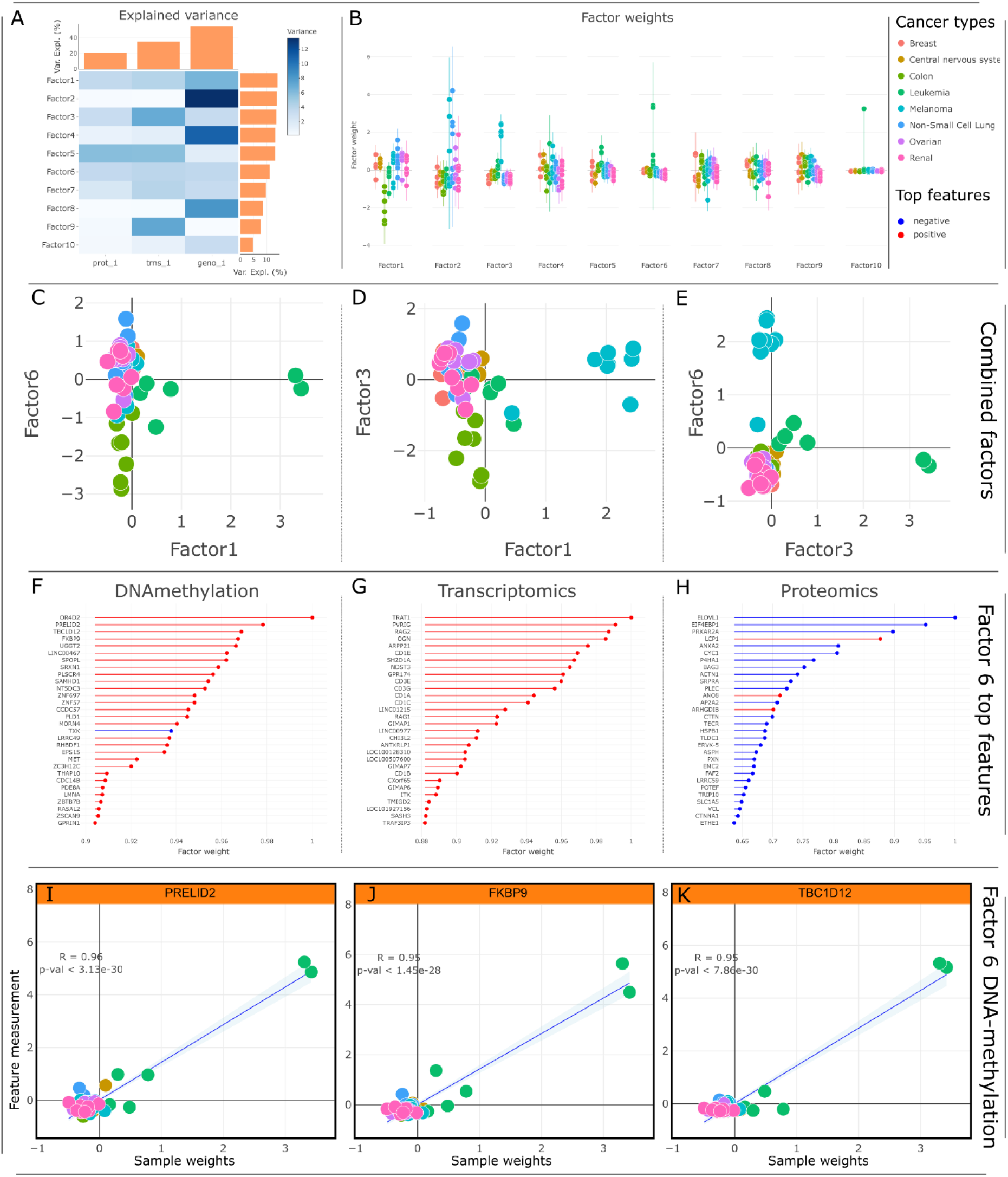
MOFA on DNA-methylation, transcriptomics, and proteomics datasets of the NCI-60 cell lines. A: explained variance for each factor and omics combinations. B: factor plot comparing sample weights stratified according to cancer type. C-E: combined factor plots showing how specific groups of interest can be discriminated using a combination of factors; in this case colon, leukemia, and melanoma show good discrimination combining factors 1, 3, and 6. F-G show top discriminating factors according to MOFA in the three omics datasets used. I-K are scatter plots for these top contributing features, plotting the actual measured value for the feature (normalized) against the sample weights.

## Discussion

In this work we introduced iSODA (integrated Simple Omics Data Analysis) for interactive single- and multi-omics data exploration. The web-based platform allows users to analyze their single-omics experiments individually, and side by side using multi-omics integration, within the same interface. This enables a more efficient streamlined analysis for multi-omics characterization experiments. Although we have demonstrated the tool in this work using two example datasets and various figures, the actual emphasis of iSODA lies in its interactive visualization capabilities, for which the true experience cannot be reflected in static figures. The modular structure of the application allows extending the core single-omics module to the specifics of the data. For example, handling lipid shorthand IDs (52). iSODA’s modularity was employed to produce SODAlight, a lipidomics-only instance designed to explore the lipidomics data of the Neurolipid Atlas project (53).

Before designing iSODA, we performed a comprehensive literature search to identify tools for single and integrative omics data analysis (Table S1), six notable software implementations stand out, MetaboAnalyst (17), Perseus (18), PaintOmics 4 (19), MergeOmics (22), OmicsAnalyst (10), and 3Omics (14) (Table 1). These software tools provide a graphical user interface (GUI), are general purpose and not specific towards a disease or an organism. We compared the software tools based on five criteria: data filtering, interactivity, functional analyses, single-omics analysis, multi-omics integration, as well as local and online interface. For **data filtering** PaintOmics, MergeOmics, 3Omics or OmicsAnalyst do not provide filtering options, requiring users to perform filtering and data preparation externally. MetaboAnalyst, Perseus and iSODA, offer advanced filtering. In regards to **interactive visualization**, MetaboAnalyst, Perseus and iSODA provide interactive plots, however this interactivity is central in iSODA allowing various plotting options as well as downloading all plots as vector graphics. All seven evaluated software packages offer some form of **functional analysis**. iSODA allows users to also supply their own annotations. Regarding providing a unified interface to **single-omics analysis** as well as **multi-omics integration**, most omics tools including the ones discussed here can be split into two categories: those specializing in single-omics (e.g. MetaboAnalyst, Perseus) and those processing multi-omics (e.g. PaintOmics, MergeOmics, 3Omics, OmicsAnalyst). iSODA provides a unified interface to perform both.

**Table 1.**
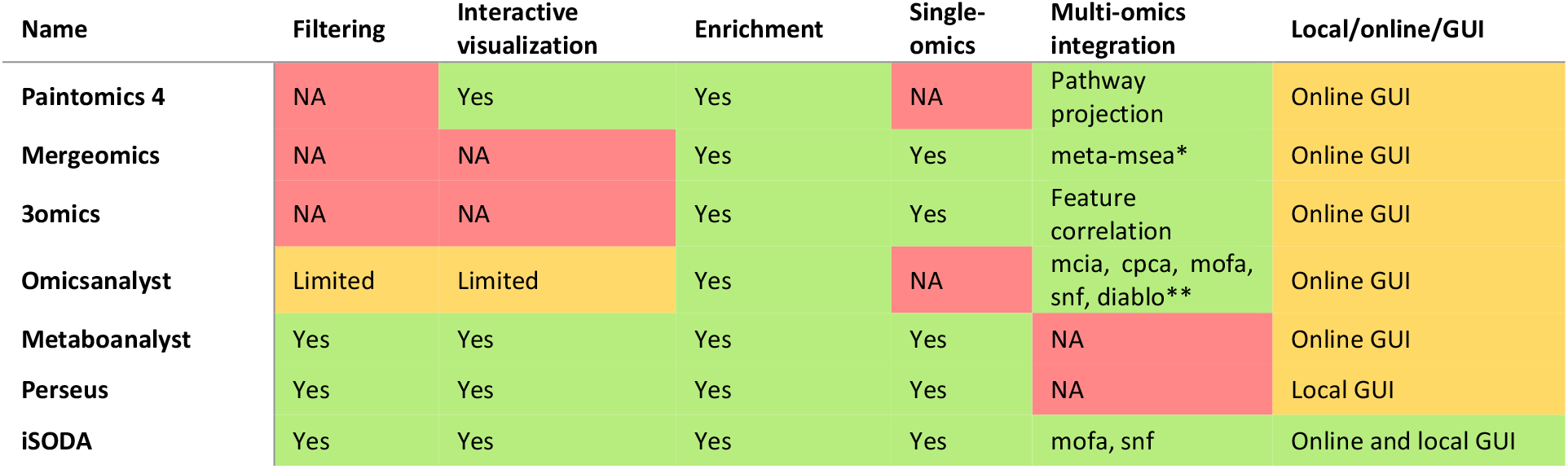
Comparison of 6 software packages: PaintOmics 4, MergeOmics, OmicsAnalyst, 3Omics, MetaboAnalyst Perseus, and iSODA. The criteria examined encompass data filtering, single-omics, interactive plots, enrichment and multi-omics integration. *Meta-MSEA: meta-Marker Set Enrichment Analysis. **Many: MCIA (Multiple co-inertia analysis), CPCA (Consensus PCA), MOFA, SNF, and DIABLO.

For the multi-omics integration, a variety of methods are used, ranging from knowledge-driven approaches based on enrichment, to data-driven approaches based on variance or similarity. Knowledge-driven methods like pathway projection (PaintOmics 4) or meta-MSEA (MergeOmics) are less effective for small molecules omics such as lipidomics. OmicsAnalyst offers the most extensive range of data-driven methods, including DIABLO and PLS-DA (supervised) and SNF, MOFA, CPCA (unsupervised). iSODA currently focuses on two unsupervised methods, SNF and MOFA. Additionally, given the unified interface of iSODA, various method for supervised feature selection (like discriminant regression analysis or statistical tests) can be performed in each single-omics modules individually, and the results carried over to the multi-omics modules. This effectively results in a supervised MOFA or SNF and shows the versatility of iSODA.

## Conclusion

The development of iSODA was driven by the necessity for a user-friendly data analysis tool that equally focusses on single-omics and multi-omics. For single-omics analysis, iSODA includes a core module with several omics-agnostic processes that are extendable to handle omics-specific analyses and visualizations. For multi-omics analysis, the goal was not only to incorporate multi-omics modules but also to connect them with the single-omics modules, leveraging the insights gained from single-omics processing to improve the integrative analysis. Importantly, iSODA is not designed as a linear analysis pipeline, it is rather a versatile toolbox allowing users to explore their data in innovative ways; going back and forth between plots and omics views, filtering and exploring specific parts of the uploaded data, all with the possibility to download the plot-associated data for external analysis if needed. Currently, the application includes five modules: lipidomics, metabolomics, proteomics, transcriptomics, and genomics. Future developments aim to introduce more specialized processes for each of these omics, as well as additional (multi-)omics modules based on user requirements.

## Acknowledgments

This work was supported by the Chan Zuckerberg Initiative, “OMICSER” grant 32620 to MG and YM.

## Data and code availability

All data used in this work is available from the software tool website at http://isoda.online as well as on GitHub https://github.com/isodaonline/iSODA.

The iSODA source code is available under GPL-3.0 license and is maintained on GitHub https://github.com/isodaonline/iSODA. The iSODA version used for this current work is also available from FigShare https://doi.org/10.6084/m9.figshare.26485198.v1.

