## supplementary materials for "iSODA: A Comprehensive Tool for Integrative Omics Data Analysis in Single- and Multi-Omics Experiments"

**Corresponding:**

Yassene Mohammed

Center for Proteomics and Metabolomics, Leiden University Medical Center, 2333ZA Leiden, Netherlands

**iSODA – integrated Simple Omics Data Analysis**

iSODA (integrated Simple Omics Data Analysis) is a software developed to simplify both single- and multi-omics data analysis.

The user starts by creating one or multiple single-omics experiments (lipidomics, metabolomics, proteomics, transcriptomics, genomics) that can be accessed individually through the side bar. Each has their own dedicated UI with three main tabs for data upload, visualization, and functional analysis respectively. After having uploaded their data, interpreted it through the visualization and looked for the underlying biological processes at play via the functional analysis, several of these single-omics modules can be imported in the multi-omics modules.

Likewise, the multi-omics modules present an upload tab to select the single-omics experiments previously created, as well as a visualization tab to interpret the results. Some biological phenomena cannot be fully understood when glancing at single-omics data, but can unravel when considering the bigger picture. MOFA can achieve that by providing more information from the features' contribution, whereas SNF focuses more or the effects of integration on the samples.

**Data tab.**

The Data tab within each single-omics module allows the user to upload their data via three subtabs named after the data they are expected to receive: Sample annotations, Measurement data, Feature annotations. While the app mainly uses the measurement data in plots and calculations, the sample and feature tables are used to annotation groups either on the samples or features, used for filtering purposes or mapped in the graphical representations of the data.

*Sample annotations:* tab designed for the upload and curation of sample metadata; the input table formatted with samples as rows and annotations as columns. This sample annotations table carries four essential columns: an ID column with unique identifiers, a type column specifying the type of each row (sample, blank, pool, or QC), a group column identifying the main sample groups to examine, and a batch column.

*Measurement data:* tab designed for the upload and curation of quantitative measurement data. The data must be provided with samples as rows, features as columns, and the values can only be numerical or missing. Upon upload, the table undergoes several normalization processes to produce other tables, for instance, total normalization and z-score scaling. Using the sample annotations, this tab can also filter the data to keep the most relevant features, displaying signals significantly above blanks. The lipidomics module expects features to be specifically formatted using lipid shorthand notations to extract structural information that is directly exported as feature annotations.

*Feature annotations:* tab used to upload feature metadata. This table is formatted with features as rows and annotations as columns. Feature annotations can be simple – one value per cell – or complex: multiple values per cell, separated by a pipe ("|") character. Complex annotations typically include ontologies, like classifications, pathways or gene ontologies and can be used in certain plots like the volcano plot, or for functional analysis.

**Interactive visualization tab.**

Interpreting results from experiments relies on processing the measured data in a specific way, depending on what needs to be achieved, and using sample and feature annotations to further enhance interpretability. The output then needs to be presented graphically in a meaningful manner for the user to assess the results and explore their data. The visualization tab is designed to automate all these steps in a few clicks: the user can select up to four plots to be displayed simultaneously and they will immediately appear on screen using the most relevant input parameters. In addition to the selection, zooming and hovering capabilities provided by plotly, iSODA's plots can be interacted with using the four sets of parameters accessed via the sidebars:

- Input settings let the user select which transformed version of the measured data to use (raw, total normalized, z-scored) as well as other relevant input to be mapped on the plot, like sample or feature annotations.
- Data settings are used to access the parameters used in statistics and other background processes.
- Aesthetics settings control the colors, marker size, opacity and other display options used to enhance the readability of the data.
- Output settings allow a selection of the output formats for images (JPG, PNG, WEBP and SVG) as well as export options within or outside the app. Some of the plot results can be exported to the annotation tables to be used elsewhere on the app, like for instance sample or feature clusters. Most plots also have associated tables which can be downloaded from here, to be used outside the app.

Networks also have their own settings to adjust the physics parameters and display clear and uncluttered networks.

Current plot selection includes a dendrogram, volcano plot, sample correlation, feature correlation, heatmap and PCA. Moreover, the lipidomics module has three specific plots: class comparison, class distribution and double bonds plots

*Dendrogram:* Provides a rapid assessment of the similarity between samples and sample groups. The samples are clustered using hierarchical clustering and displayed in the form of a dendrogram, which can be mapped with sample annotations to find out if the unsupervised groups can be explained by known factors. The number of desired clusters can be set and saved on the sample annotations table to be used elsewhere on the app. For example, in Figure 4 one can see how this functionality is used to highlight that cancer cell lines are primarily clustered according to whether the cells are epithelial or not.

*Volcano Plot:* A standard way of identifying features that distinguish two sample groups. The features are displayed in a scatter plot, the y-axis representing the -Log10(p-value) and the x-axis the Log2(fold change). The p-values are calculated using either a t-Test or a Wilcoxon test and can be adjusted with multiple methods. The fold changes are calculated using either the mean or the median values. In some cases, p-values or fold changes cannot be calculated because of one group having too few or no values for a feature. These features are reported in violin plots surrounding the main scatter plot. The left-to-right spread of the features reflects their differential production or expression between the two sample groups (higher absolute fold change), and the more a feature is located on the top of the plot, the more significant it is (lower p-value). The adjacent violin plots display the features that can be regarded as some of the most important, since they are absent in one of both groups. To highlight relevant features, the user can set p-value and fold change thresholds, manifesting in dashed lines across the volcano plot. Features above these thresholds can be exported to the feature annotations table and used, for instance, in functional analysis. As an alternative to viewing the differential expression, the user can also map feature annotations on the volcano plot to assess if a known group of features is being over- or underexpressed.

*Sample Correlation Plot:* An alternative representation to the dendrogram visualization, augmented with a heatmap to display and better understand sample clusters. The heatmap displays the correlation coefficient (Pearson or Spearman) between each sample pair, and the sample order is arranged using hierarchical clustering, the results of which are also displayed with the dendrograms on the sides of the heatmap. As with the dendrogram, sample annotations can be mapped on the leaves of the dendrograms. While the dendrogram only displayed the clusters, the sample correlation heatmap also shows how close (or different) samples are based on the correlation coefficients. The sample clusters are also emphasized with the heatmap colors. The calculated clusters can be exported to the sample annotations table.

*Feature Correlation Plot:* Shows which features are correlated and highlights feature groups thanks to the hierarchical clustering and a heatmap. It is the same representation as the sample correlation but applied to features. Due to the limited rendering capabilities on browsers, the plotted data can be reduced using a maximum feature count filter and a minimum correlation coefficient threshold, thus displaying only the best scoring correlations. Like the sample correlation, the correlation coefficients are shown on the heatmap, and the feature ordering is based on hierarchical clustering, helping highlight groups of features and their correlations, notably with the color coding. Obtaining feature annotations is not as straightforward as gathering sample information: creating groups of features based on this feature correlation is a good starting point when nothing else is available. The generated feature clusters can then be exported to the feature annotations table.

*Heatmap:* A visual representation of the measured data combining sample and feature clustering, allowing to find out if some groups of samples are associated with groups of features. This is achieved by displaying the z-score scaled data as a heatmap with samples as columns and features as rows. Based on the data, hierarchical clustering is applied to samples and features, highlighting areas in the dataset where groups of samples and feature expression coincide. Since this plot combines sample and features, their respective annotations can be mapped on the heatmap. Moreover, Lasso and Elastic-Net Regularized Generalized Linear Models can be applied on the features to keep only those best segregating the sample groups. Using the representation, the user can directly assess if the sample group mapping can be associated with known feature groups, driving the observed clustering.

*Principal Component Analysis:* Representation of the samples and the features in a reduced dimensional space that retains as much variance as possible from the dataset, highlighting potential sample groups and the features driving the groupings. In iSODA, PCA is delineated in three plots: explained variance, scores plot and loadings plot, with the option to display the scores and loadings plots together. The user can choose the number of computed principal components and display the associated explained variance. Two PCs can then be chosen for the 2D scores and loadings plots. The scores plot displays the coordinates of the original samples in this new 2D space. Likewise, the loadings plot represents each feature's contribution to the two selected PCs. Sample and feature annotations can be mapped on the markers. The scores plot can be used to identify trends and sample groupings, while the loadings plot can identify the features contributing to these trends. Like with the heatmap, Lasso and Elastic-Net Regularized Generalized Linear Models can be used to specifically select the most segregating features.

*Class Distribution:* A lipidomics specific visualization that provides a summary of the mean lipid class concentrations for all sample groups. The lipid concentration is displayed on the y-axis and the lipid classes on the x-axis. The class-wise group concentrations will be represented as colored bar plots. This plot allows the user to directly spot concentration differences between sample groups and assess the relative lipid class concentrations with the shared y-axis.

*Class Comparison:* Grid version of the class distribution, allowing a better assessment of the minute group concentration variations. Sample group concentrations are represented by bar plots for each lipid class. Plots are arranged in a grid, each one with its own y-axis. Like for the class distribution, the bars represent the mean concentrations. On top of the bars, box plots represent the median concentrations and the quartiles, along with the individual sample concentrations as markers. With the separate axes, group concentrations can be better examined. The additional box plots and markers help identify the sample distribution for each group and potential outliers.

*Double Bonds Plot:* Another lipidomics specific plot that works in conjunction with the volcano plot, allowing a more structure specific examination of the lipid class differences between two sample groups. Two sample groups and a lipid class are selected, and like with the volcano plot, p-values and fold changes are calculated. Each individual lipid species is displayed on a bubble plot, the y-axis representing the double bond count and the x-axis the carbon count. The carbon and double bond counts can be specified to the side-chain level or the total values. Each bubble – or lipid species – is displayed with a size relative to the p-value and colored according to fold change. The bubble size emphasizes the most relevant features while the coloring highlights the direction of the expression. Combined with the structural information provided by the double bonds and carbon counts, this plot can reveal structural trends that might be biologically relevant to differentiate the two conditions.

**Functional analysis tab.**

This tab is designed for functional analysis, a deeper dive to reveal the factors differentiating two sample groups. It is subdivided into three subtabs, “Functional comparison” to set up the analyses, and two visualization tabs to plot the results: “Enrichment” and “Over-representation”. The functional comparison tab is divided into multiple sections to prepare the input data and the parameters for the analyses. The sample selection lets the user choose the appropriate data table and the two sample groups to compare. The feature selection section lets the user choose which features will be kept for the subsequent functional analysis. This can be done either with a statistical in conjunction with p-value and fold change thresholds; or using a custom user selection via the feature annotation table, which includes groups exported from other plots, like the volcano plot and the feature correlation. The last two sections are dedicated to the parameters for enrichment (EA) and overrepresentation (ORA) analyses, notably selecting the “Feature sets” to be used. Feature sets are groups of features sharing a role, like the Gene Ontology “Biological Processes”, “Molecular Functions” and “Cellular Components”, which are automatically available for non-small molecule omics. Alternatively, the feature sets can be supplied in the feature annotation table. Since each feature can be associated with multiple feature sets, these can be separated by a pipe (“|”) character.

***Enrichment analysis.*** Computational method designed to determine whether feature sets show statistically significant, concordant differences between two sample groups. It is based on the original Geneset Enrichment Analysis (GSEA) but applied to any omics using appropriate feature sets. The algorithm starts by ranking all features in the dataset based on their differential expression between the two sample groups, creating a ranked list. An Enrichment Score (ES) is then calculated for each predefined feature set, which reflects the degree to which features from that set are represented at the extremes of the ranked feature list. This is achieved by walking down the ranked list and increasing a running-sum statistic when a feature in the feature set is encountered and decreasing it when it is not. The ES is the maximum deviation from zero encountered in walking the list, corresponding to a weighted Kolmogorov-Smirnov-like statistic. A positive ES indicates a feature set enrichment at the top of the ranked list while a negative ES indicates a feature set enrichment at the bottom of the ranked list. Subsequently, a permutation test is applied to estimate the significance of the ES and derive a p-value, indicating the likelihood that the observed enrichment is due to chance. The permutations are then used to calculate the Normalized Enrichment Score (NES) to account for differing set sizes. By applying a feature set p-value threshold, the enrichment analysis can identify sets that are significantly associated with one sample group or the other. Within these sets, some features contribute more than others to the enrichment score: the core enrichment – or leading-edge subset – are features that appear in the ranked list at or before the point where the running sum reaches its maximum deviation from zero.

***Over-representation analysis (hypergeometric test / fisher exact test).*** Over-representation Analysis (ORA) is a statistical method used to identify if a feature set is represented more frequently in a list of pre-selected features of interest than would be expected by random chance. These pre-selected features of interest are set apart from the other features either by using a statistical test along with p-value and fold change thresholds; or selected manually by the user via the feature annotations table. A hypergeometric test (or Fisher's exact test) is then applied to assess the probability (p-value) that the observed frequency of a specific feature set within the list occurs more than would be expected by chance, given the distribution of annotations in the universe of features (i.e. the complete list of features). Sets with a low p-value are significantly over-represented in the list of features of interest, implying they are relevant in the studied sample groups.

**Functional analysis plots.** Enrichment and over-representation analysis use similar plots that rely on the same metrics, highlighting the top feature sets and features that differentiate the compared sample groups.

*Dot Plot:* This plot displays the most significant feature sets resulting from the analysis. They are represented on the y-axis of the bubble plot, against their associated feature ratio on the x-axis. Each bubble – or set – is colored based on their p-value and sized proportional to their feature count. For enrichment analysis, the plot differentiates between suppressed and activated sets using enrichment scores, highlighting the sets associated with one sample group or the other. This visualization aids in identifying biologically significant feature sets that contribute to the observed differences between the two groups under comparison.

*Bar Plot:* This plot illustrates the top feature sets identified from an overrepresentation analysis, similar to the previously described dot plot. The y-axis lists the feature sets, while the x-axis shows the feature ratios. Each bar is colored according to the p-value of the set it represents, showcasing a clear visual comparison of the statistical significance across different feature sets.

*Ridge Plot:* This ridge plot provides a multi-layered visualization of the top feature sets identified in an enrichment analysis, with each set plotted along the y-axis against the Log2(fold change) of its associated features on the x-axis. The distribution of features within a set is represented by a density curve along the x-axis, creating a series of vertically stacked ridges. Each ridge is color-coded according to the p-value of its corresponding feature set. Horizontally, the plot illustrates the spread of features from one extreme to the other, indicating their predominance in one of the sample groups. Vertically, the plot allows for comparison of these feature distributions across different sets.

*CNET Plot:* This network visualization represents feature sets and their associated features as nodes, connected by edges that link features to their respective feature sets. Annotations for both features and sets can be mapped onto the nodes to provide additional information. The CNET plot provides a way to visualize and interpret the complex relationships between features and their feature sets. By displaying Log2(fold change) values on the feature nodes, the plot facilitates the observation of potential differential expressions among feature sets. The plot not only shows which feature sets are influenced by the current analysis but also highlights the associated features. Since these features may belong to multiple sets, the plot often reveals clusters of feature sets, indicating broader, more complex phenomena that aggregate the effects of individual sets.

*eMap Plot:* The enrichment map (eMap) plot is a streamlined alternative to the CNET plot, specifically designed to handle large numbers of feature sets without becoming cluttered. Unlike the CNET plot, the eMap plot omits feature nodes, reducing complexity and avoiding the creation of unreadable "hairball" networks. Instead, it creates set-to-set edges that reflect the number of shared features relative to the total associated features, quantified by Jaccard's similarity score (default) to indicate connection strength between two feature sets. The network can be further simplified by applying a similarity score threshold to produce more manageable feature set clusters. Node sizes are scaled based on the feature count of each set, while node coloring reflects p-values, and edge thickness is proportional to the similarity score. This layout allows users to discern larger biological mechanisms coalescing smaller feature sets, offering insights on a bigger scale than what is provided by the CNET plot.

**Multi-omics integration**

The main aim of omics integration is finding and possibly quantifying patterns in the multiple omics measurements that are not obvious when considering them separately. Each omics layer (genomics, proteomics, transcriptomics, metabolomics) provides a different perspective on the biological system, their integration offers a holistic view of the biological processes and mechanisms at play.

***MOFA (Multi-Omics Factor Analysis).*** MOFA is an unsupervised integration method designed to reduce the complexity of large-scale omics datasets into a manageable number of latent factors (Z). These factors capture the underlying sources of variation across the datasets, providing insights into biological processes that might be driving the observed patterns. Factors can be assimilated to principal components in PCA, each explaining a portion of the variance from the dataset. Similarly, samples and features have weights (W) associated to each factor.

*Explained Variance Plot:* This plot provides an overview of the contribution of each omics dataset to the computed factors by displaying the variance explained for each factor in the form of a heatmap. Additionally, two bar plots on the sides show the cumulative variance for each omics and each factor. This arrangement allows researchers to identify factors unique to specific omics as well as those that are significant contributors across multiple omics, potentially indicating a shared biological phenomenon. The plot serves as a useful reference in conjunction with other plots to identify the most relevant factors for the omics under investigation.

*Factor Plot:* This plot summarizes the sample factor weights and assesses their potential to explain group differences. It displays samples on the x-axis and their corresponding factor weights on the y-axis. Multiple factors can be represented simultaneously, with samples differentiated according to sample annotations. Optional violin plots give a better impression of sample distributions. By mapping sample annotations, researchers can determine whether certain factors distinctly separate specific sample groups. These factors may then be analyzed in greater detail using additional visualizations.

*Combined Factors Plot:* This visualization complements the factor plot by showing whether paired factors explain variations between sample groups. One or multiple factors can be selected, which generates a grid containing two types of plots. Plots on the diagonal illustrate the distribution of samples across the loadings of a single factor, while off-diagonal plots depict the distribution across two factors. Density plots are used to visualize group distributions and to assess whether factors explain differences between sample groups. Additionally, the combined factor scatter plots enable the examination of whether sample groups can be characterized by more than one factor, similar to how PCA plots sample scores for two components.

*Feature Weights Plot:* This plot illustrates the contributions of individual features to a selected factor within a specified omics dataset. Users select a factor and omics dataset, and the features are displayed with their factor weights on the x-axis and their contribution rank on the y-axis. Typically, features will align along a sigmoid curve, or alternatively, a logarithmic curve when absolute factor weights are considered. This format allows users to quickly identify the most influential features at either end of the sigmoid curve or at the peak of the logarithmic curve. Additionally, mapping feature annotations can provide deeper insights into which groups of features significantly influence the factor.

*Feature Top Weights Plot:* This plot serves as an alternative to the feature weights plot, focusing specifically on the highest-scoring features. Users select an omics-factor pair, and the plot displays the top contributing features in a lollipop plot. Contributions can be further filtered to show either the top negative or positive contributing features, enabling a more targeted analysis of feature impact.

*MOFA Heatmap:* This visualization integrates the single-omics heatmap approach with the identification of a factor's top contributing features. Similar to the heatmap available in the single omics module, the MOFA heatmap presents the measured data and provides clustering for both samples and features. Unlike the standard heatmap, which employs supervised discriminant analysis for feature selection, the MOFA heatmap selects the top-ranked features based on their factor contributions, regardless of whether these are negative or positive. This enables the user to view if certain sample groups can be distinguished by the most relevant features of a factor. If no distinct groups are apparent, it may indicate that the factor is associated with an overlooked biological process.

*Scatter Plot:* This plot enables users to assess the correlation between specific feature signals and the dataset's sample weights. Users can select an omics dataset along with the top contributing features to a factor—whether negative, positive, or both. A grid of scatter plots is then generated, each representing a distinct feature. In each scatter plot, samples are plotted with their measured values for that feature on the y-axis and their sample weights on the x-axis. A Pearson correlation coefficient and a confidence interval are calculated and displayed for each plot to quantify the strength and certainty of the correlation. Sample annotations can be mapped to visually emphasize different sample groups. This scatter plot provides a detailed examination of how the measured top-weighted features correlate with sample weights, possibly explaining sample groups.

***SNF (Similarity Network Fusion).*** SNF is another computational method for integrating different single-omics datasets. SNF works by constructing individual networks for each dataset, each representing the similarity between samples. These networks are then fused into a single network through an iterative process capturing both the shared and unique characteristics of each dataset. For a single-omics dataset, a distance matrix between the samples is created, using the distance method chosen by the user (e.g. Euclidean). The distance matrix is then transformed into an affinity matrix, capturing the local neighborhood relationships among the samples by employing the the K-nearest neighbors and sigma parameters. The former specifies the number of nearest neighbors for each sample based on the distance matrix, and the latter determines how rapidly the affinity between samples decreases when constructing the affinity matrix. The samples are then grouped into K clusters through spectral clustering. In the case of the fusion heatmap and the similarity fusion network, the affinity matrices of each individual omics are combined, using the additional K-nearest neighbors parameter and a designated number of iterations (T) for the diffusion process, thereby enhancing the comprehensive understanding of the dataset through a multifaceted approach.

*Similarity Heatmap:* This heatmap illustrates the impact of spectral clustering on omics datasets separately. The x-axis and y-axis display samples organized by cluster groups, while the cell values indicate affinity levels. Cluster assignments can be shown as side annotations alongside other sample characteristics, enabling evaluation of whether the clustering within single-omics data aligns with anticipated sample groupings.

*Fusion Heatmap:* This heatmap presents sample affinities by integrating multiple omics datasets. Users can select two or more single-omics datasets, which are each subjected to spectral clustering. The outcomes are then merged to form a comprehensive affinity matrix that encapsulates the characteristics of the integrated omics data. Like the similarity heatmap, this heatmap organizes samples according to their cluster affiliations and allows for the mapping of sample annotations. The fusion heatmap is particularly useful for identifying sample similarities that emerge when multiple data perspectives are considered simultaneously.

*Similarity Network:* This network graph visualizes the affinity matrix for single-omics datasets. The similarity matrix is converted into a network, where all samples are connected by edges that denote the affinity values between them. To enhance readability, the network is trimmed to retain only the highest-scoring edges, a percentage which can be set by the user. Nodes are color-coded based on cluster affiliation or other sample annotations, and edge thickness is proportional to the affinity value. By interactively hovering over nodes and edges, additional details will be displayed on screen. This network serves as an alternative to the similarity heatmap, enabling users to verify whether the clustering of single-omics data corresponds with predefined sample groups and to explore complex relationships like affinities across multiple sample groups.

*Similarity Fusion Network:* This network graph represents the multi-omics sample clustering, employing the same approach as the single-omics similarity network but based on the fusion heatmap. Users can select two or more single-omics datasets, and the combined affinity matrix is then transformed into a network. Here again, the user can select the top scoring edges to be displayed. The edge width corresponds to the affinity values, while the coloring denotes the number of omics datasets supporting each connection, based on the set edge threshold. This network is interpreted like the single-omics network, with the added benefit of sourcing the connections from multiple datasets. The similarity fusion network is particularly effective in revealing groupings less apparent in single-omics analyses. It also identifies unique connections that emerge only when multiple omics datasets are considered together.

### Supplementary Figures and Tables

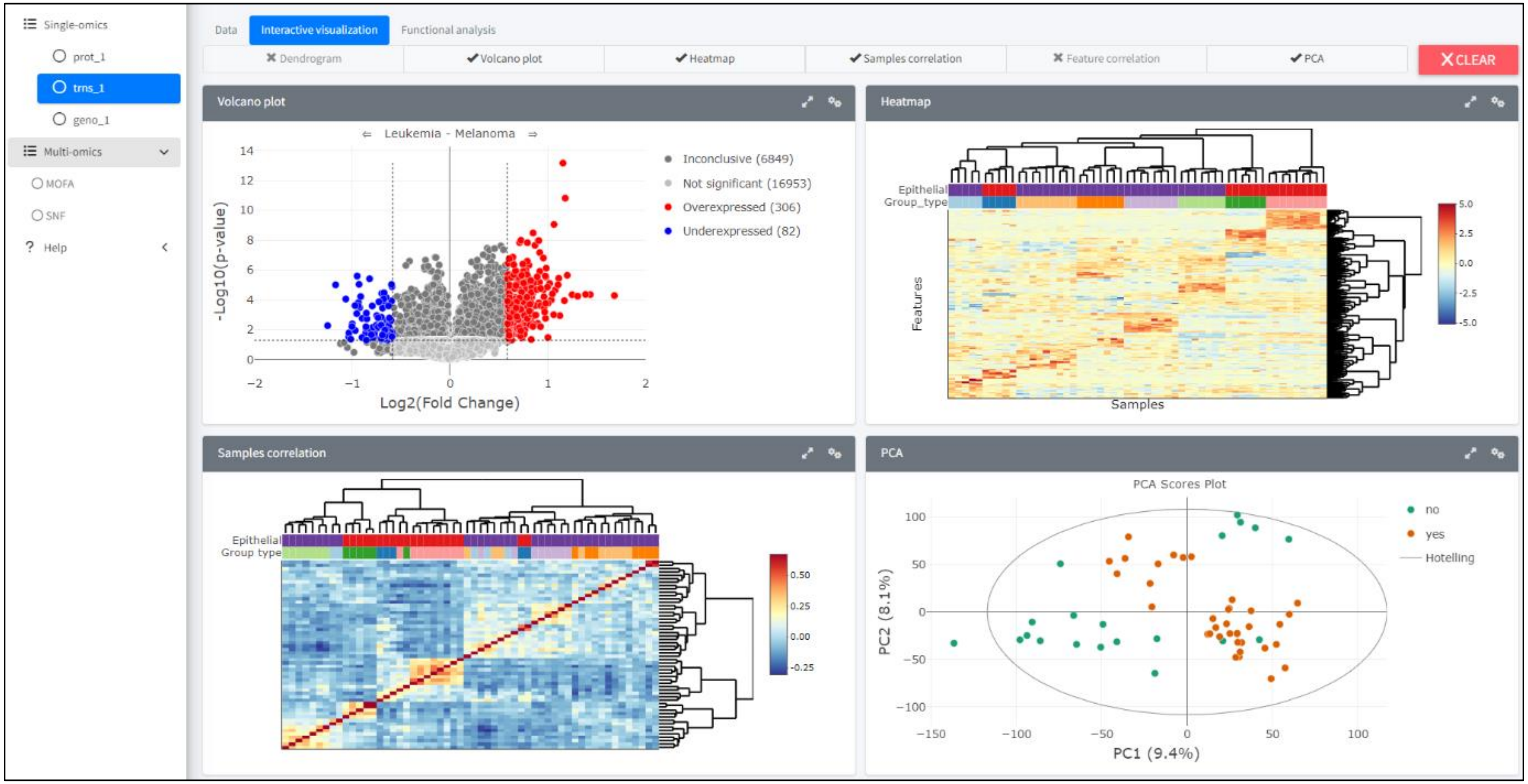


Figure S1. Screenshot of iSODA showing four interactive analyses

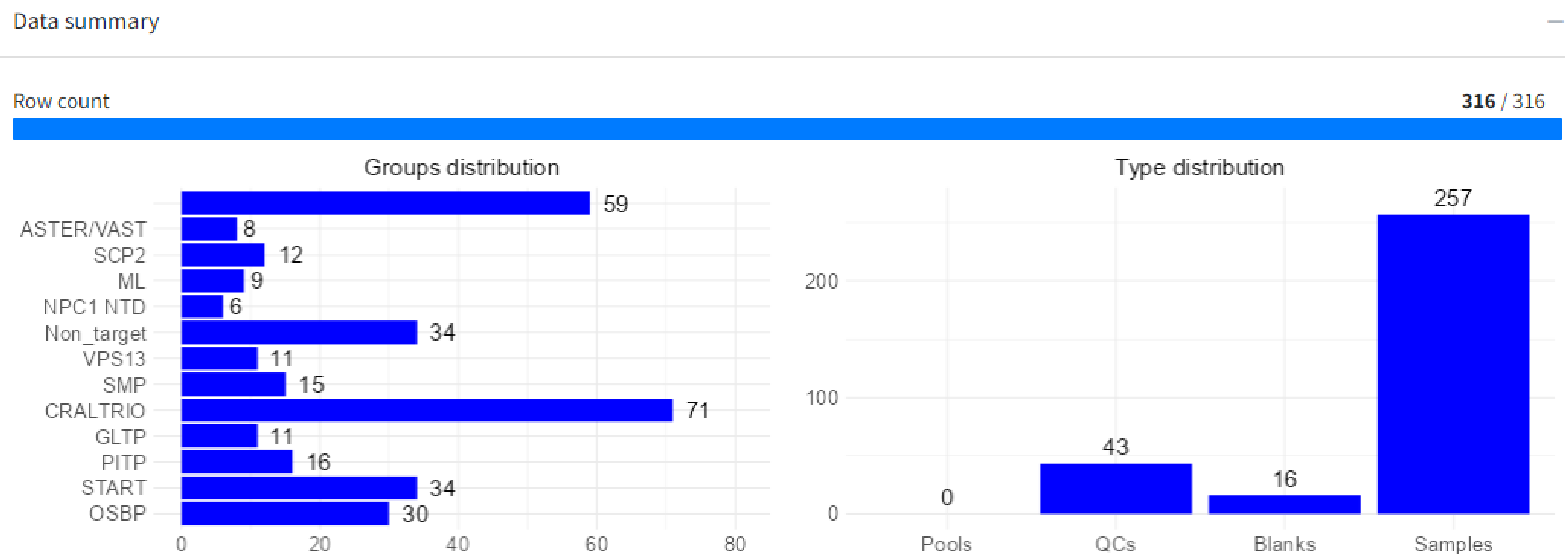


Figure S2. A screenshot of the data summary of the LTP knockout library dataset as shown in iSODA.

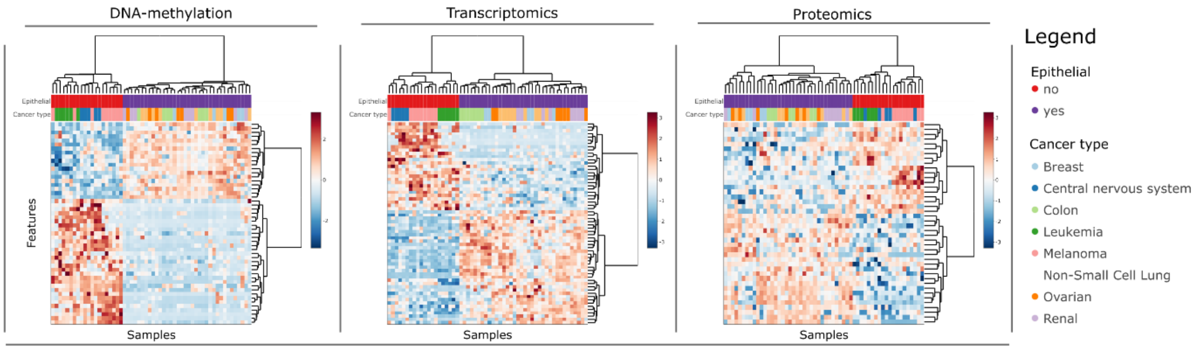


**Figure S3.** Heatmaps with feature selection to discriminate epithelial nature for the three omics (z-scored total normalized data, LASSO alpha set to 0.8, Euclidean distance, ward.D2 clustering). Annotations mapped on top included cancer type and epithelial nature of the tissue. k=8 clusters were set and rendered in the colored dendrogram.

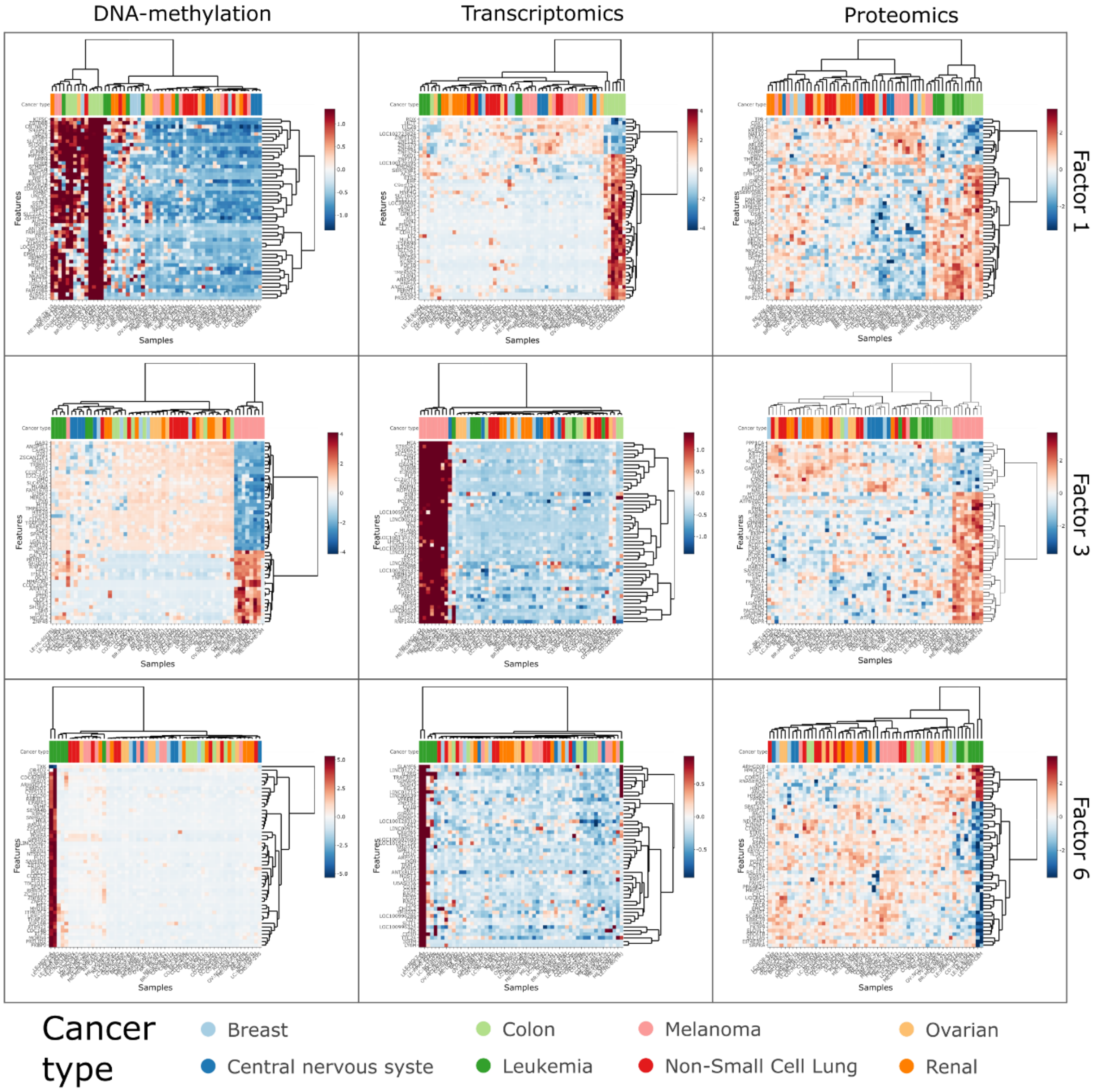


**Figure S4.** MOFA heatmaps for factors1, 3 and 6 across all datasets, displaying the top 50 contributing features (positive or negative contributions.

Table S1: tools researched for omics data analysis.

| Name | Year | Theme | URL |
|---|---|---|---|
| *KaPPA-View* | 2005 | Plants | https://doi.org/10.1104/pp.105.060525 |
| *g:Profiler* | 2007 | Gene data | https://doi.org/10.1093/nar/gkm226 |
| *MetPA* | 2010 | Metabolomics data | https://doi.org/10.1093/bioinformatics/btq418 |
| *Paintomics* | 2010 | General | https://doi.org/10.1093/bioinformatics/btq594 |
| *IMPaLA* | 2011 | General | https://doi.org/10.1093/bioinformatics/btr499 |
| *Metscape 2* | 2011 | General | https://doi.org/10.1093/bioinformatics/btr661 |
| *VisANT 4.0* | 2013 | Networks | https://doi.org/10.1093/nar/gkt401 |
| *3Omics* | 2013 | General | https://doi.org/10.1186/1752-0509-7-64 |
| *CARMO* | 2015 | Plants | https://doi.org/10.1111/tpj.12894 |
| *Mergeomics* | 2016 | General | https://doi.org/10.1186/s12864-016-3057-8 |
| *OASIS* | 2016 | Cancer | https://doi.org/10.1038/nmeth.3692 |
| *Pathview Web* | 2017 | General | https://doi.org/10.1093/nar/gkx372 |
| *OmicsNet* | 2018 | Networks | https://doi.org/10.1093/nar/gky510 |
| *PaintOmics 3* | 2018 | General | https://doi.org/10.1093/nar/gky466 |
| *miRNet 2.0* | 2020 | miRNA | https://doi.org/10.1093/nar/gkaa467 |
| *Argonaut* | 2020 | General | https://doi.org/10.1016/j.patter.2020.100122 |
| *Imaging-AMARETTO* | 2020 | Imaging | https://doi.org/10.1200/CCI.19.00125 |
| *CHOmics* | 2020 | Cell lines | https://doi.org/10.1371/journal.pcbi.1008498 |
| *ADAS-viewer* | 2020 | Alzheimer | https://doi.org/10.1038/s41540-021-00177-7 |
| *Panomicon* | 2020 | Gene data | https://doi.org/10.1016/j.heliyon.2020.e04618 |
| *GraphOmics* | 2021 | General | https://doi.org/10.1186/s12859-021-04500-1 |
| *Arena3D* | 2021 | Networks | https://doi.org/10.1093/nar/gkab278 |
| *NeDRex* | 2021 | Drug repurposing | https://doi.org/10.1038/s41467-021-27138-2 |
| *Mergeomics 2.0* | 2021 | General | https://doi.org/10.1093/nar/gkab405 |
| *MiBiOmics* | 2021 | General | https://doi.org/10.1186/s12859-020-03921-8 |
| *OmicsAnalyst* | 2021 | General | https://doi.org/10.1093/nar/gkab394 |
| *TIMEOR* | 2021 | Gene data | https://doi.org/10.1093/nar/gkab384 |
| *MAINE* | 2021 | Gene data | https://doi.org/10.1093/bioinformatics/btab862 |
| *iNetModels 2.0* | 2021 | Database | https://doi.org/10.1093/nar/gkab254 |
| *MOVIS* | 2022 | General | https://doi.org/10.1016/j.csbj.2022.02.012 |
| *OmicsNet 2.0* | 2022 | Networks | https://doi.org/10.1093/nar/gkac376 |
| *MicrobioSee* | 2022 | Microbiology | https://doi.org/10.3389/fgene.2022.853612 |
| *HTT-OMNI* | 2022 | Huntingtin | https://doi.org/10.1016/j.mcpro.2022.100275 |

| | | | |
|---|---|---|---|
| *ExpressVis* | 2022 | Trns & Prot | https://doi.org/10.1093/nar/gkac399 |
| *Visual Omics* | 2022 | Gene data | https://doi.org/10.1093/bioinformatics/btac777 |
| *BALDR* | 2023 | Diabetes | https://doi.org/10.1371/journal.pcbi.1011403 |
| *PlantMetSuite* | 2023 | Plants | https://doi.org/10.3390/plants12152880 |
